# Allele-specific genomics decodes gene targets and mechanisms of the non-coding genome

**DOI:** 10.1101/2025.03.03.641135

**Authors:** Tim P. Hasenbein, Sarah Hoelzl, Stefan Engelhardt, Daniel Andergassen

**Affiliations:** Institute of Pharmacology and Toxicology, Technical University Munich (TUM), 80802, Munich, Germany; DZHK (German Centre for Cardiovascular Research), Partner Site Munich Heart Alliance, Munich, Germany

## Abstract

A large proportion of disease variants is found in non-coding RNAs (ncRNAs), gene loci that have been identified as key regulatory elements. However, for most ncRNAs, their targets are unknown, hindering our ability to understand complex diseases. Here, we found that allele-specific ncRNAs were enriched nearby allelic protein-coding genes (pcGenes), suggesting that the allele-specific information could be used to predict *cis*-acting ncRNA-targets. We translated this concept into the Allelome.LINK framework and applied it to the major mouse organs, revealing 397 events where the allele-specific expression (ASE) of a ncRNA correlated with the ASE of a nearby pcGene, suggesting either enhancing or repressive regulatory interactions. Next, we applied our strategy to the largest human dataset including tissues from nearly 1,000 individuals. Given the outbred nature of humans, each individual allows for the discovery of novel ASE correlation events. We uncovered 2,291 shared ASE events along with their mechanisms, which we benchmarked against sample-matched eQTLs, yielding a high validation rate of 77.47%. Further GWAS integration assigned 30.59% of variants overlapping informative ncRNAs to their pcGene targets. As more sequencing data and risk variants become available, this strategy has the potential to decode the entire *cis*-acting landscape of the non-coding genome.

## Introduction

Only about 1% of the human genome consists of protein-coding genes (pcGenes), leaving the majority as non-coding. These non-coding segments contain essential regulatory elements, including promoters, enhancers, and non-coding RNAs (ncRNAs), that determine the temporal and spatial activation or repression of genes. This process of transcriptional regulation is implicated in numerous human diseases, as the majority of disease-associated variants are located within non-coding regulatory regions and exert their effects through gene regulation^1,2^. While regulatory elements across tissues and developmental stages have been extensively mapped^3–5^, and genome-wide association studies (GWAS) have helped catalog human genetic disease variants^6^, the specific target genes and mechanisms-of-action remain largely elusive. This knowledge gap hinders our ability to understand how variants within non-coding regulatory regions cause disease, which is important for developing effective treatments.

Eighty percent of the human genome undergoes transcription, highlighting the prominence of ncRNAs within the non-coding genome^5,7^. Recent database analysis indicates that a significant fraction of disease-associated variants overlap with ncRNA loci, underscoring their potential significance in the context of diseases^8^. The most prominent sub-class of ncRNAs are long non-coding RNAs (lncRNAs). Over the past decade, more than 60,000 lncRNAs have been identified in various human tissues^9^, which has generated considerable interest in understanding their role in gene expression regulation. In contrast to pcGenes, ncRNAs display a remarkable degree of tissue-specificity and much greater inter-individual expression variability among humans^10,11^.

However, due to their highly dynamic expression patterns, traditional approaches, such as genotype correlation studies reliant on a wide range of biological samples, are not sufficient to identify ncRNA targets. Here, we propose a novel concept to predict the target genes of *cis*-acting ncRNAs, based on the allele-specific expression (ASE) pattern.

ASE is the favorable expression of one of the two alleles in diploid organisms, which distinguishes them from the majority of genes that show equal expression from both alleles. The vast majority of ASE is due to genetic differences between the alleles^12,13^, that disrupt transcriptional regulation by impacting the binding of transcription factors to promoters or enhancers^12,14^, interfering with the function or activity of regulatory ncRNAs, or influencing allelic gene expression post-transcriptionally by affecting splice site selection or RNA stability^15^. Genome-wide mapping of ASE has been performed in many human^16,17^ and mouse tissues^12,18^. We previously created a comprehensive survey of ASE and observed a correlation between the number of allele-specific ncRNAs and pcGenes^12^, implying a potential co-regulatory relationship.

In this study, we further explored this finding and observed a significant clustering of allele-specific ncRNAs around allele-specific pcGenes in mice and humans. Given the rarity of ASE, this suggested that the allele-specific information could be used to predict the target genes of ncRNAs. We translated this concept into a bioinformatics pipeline termed Allelome.LINK to identify correlating ncRNA-mRNA ASE events, enabling the prediction of the targets and regulatory mechanisms of *cis*-acting ncRNAs. Applying our strategy to major organs in mice and exploiting inter-individual variation in humans, we identified 397 and 2,291 ncRNA-mRNA ASE events, respectively. Further integration of non-coding GWAS variants, linked a significant proportion of non-coding disease variants to their protein-coding targets. As more individual sequencing data and risk variants become available, this strategy will decode the functional landscape of the non-coding genome.

## Results

### Allele-specific ncRNAs are strongly enriched in proximity to allele-specific pcGenes

To investigate the correlation between the number of allele-specific ncRNAs and the number of allele-specific expressed pcGenes, we first comprehensively mapped the allele-specific transcriptome of 9-week-old female F1 hybrid mice, generated by crossing BL6 females with CAST males (see Materials and methods). This analysis encompassed six primary mouse organs: brain, heart, lung, liver, kidney, and spleen (replicates *n* = 3; Figure 1A and S1). Only high-confidence loci informative in all replicates were retained for downstream analysis. Genes with an allelic ratio ≥ 0.7 or ≤ 0.3 were classified as allele-specific (Figure 1B). On average, 8.98% (*n* = 1039) of the informative genes showed ASE per tissue, ranging from 10.5% (liver, *n* = 1007) to 6.4% (brain, *n* = 840, Figure 1C, Table S1). Among these, a mean of 2.13% were ncRNAs, mainly corresponding to lncRNAs (69.4%, Figure 1D). Next, we compared the abundance of ncRNAs and pcGenes with an allelic bias across all tissues and found a positive correlation between allele-specific ncRNA proportions and allele-specific pcGenes (Spearman correlation: R = 0.66, *p* = 0.004, Figure 1E) consistent with our previous observations^12^. To investigate whether the allele-specific ncRNAs were significantly enriched near allele-specific pcGenes, suggesting that they might be co-regulated, we quantified their co-occurrence within various window sizes (Figure S2). As a control, we calculated the enrichment of allele-specific ncRNAs to biallelic pcGenes. This analysis revealed a significant enrichment of allele-specific ncRNAs around allele-specific pcGenes up to ±100kb distance (Wilcox-Test, *p* = 0.002, Figure 1F, Table S2), which indicates a co-regulatory association between them. This further suggests that the allele-specific information can be exploited to predict the *cis*-targets of ncRNAs. In addition, the underlying regulatory mechanism may be inferred based on the expression pattern: a repressive interaction when both the ncRNA and the nearby pcGenes show expression bias toward opposite alleles, and an enhancing interaction when there is a correlative bias from the same allele (Figure 1G).

**Figure 1.**
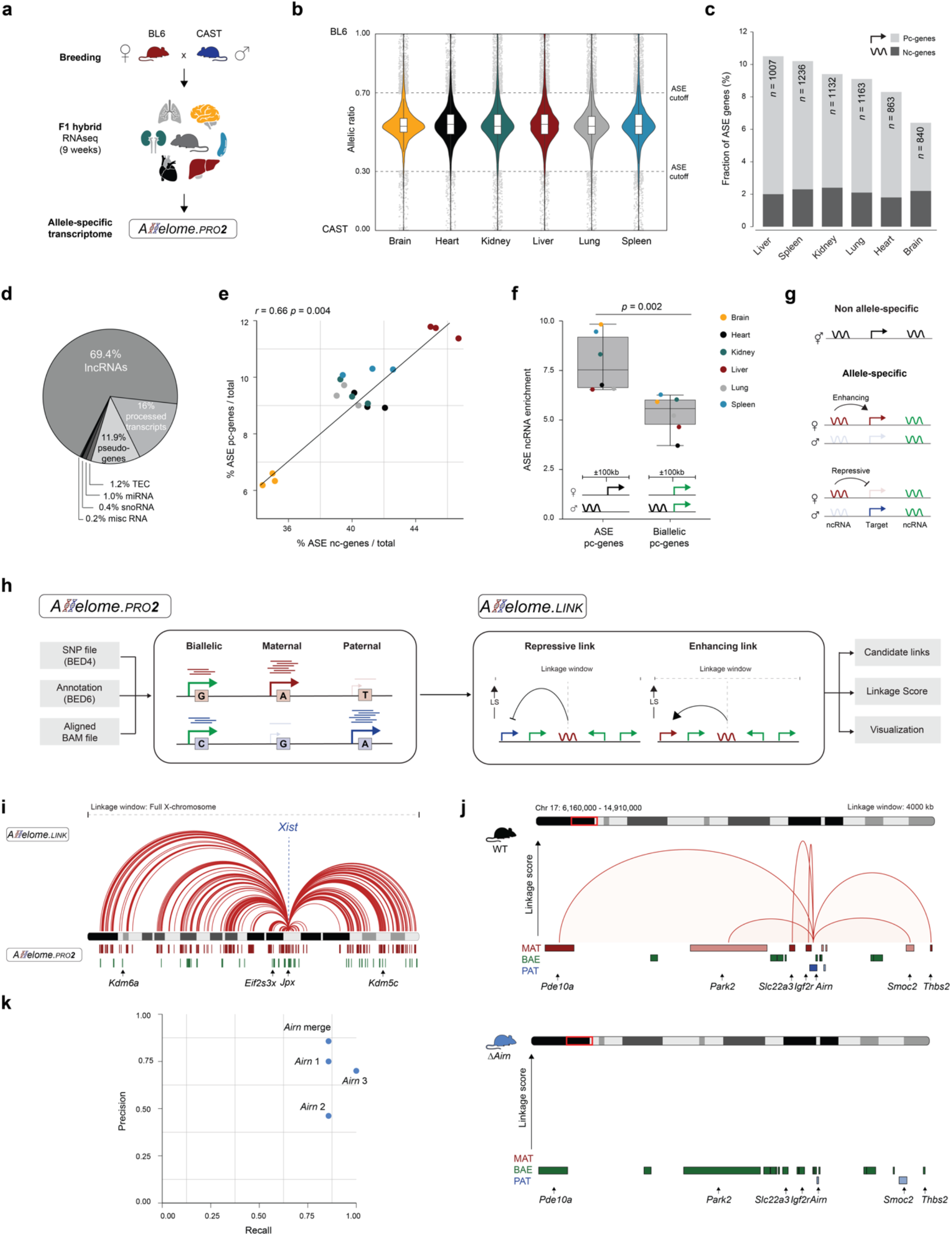
Allele-specific non-coding RNAs show enrichment in close proximity to allele-specific protein-coding genes. (**A**) Schematic overview of the allele-specific mapping process. Using Allelome.PRO2, we mapped the allele-specific transcriptome of RNA-sequencing data acquired from F1 hybrid mice (BL6 x CAST). Tissue samples were collected from the brain, heart, lung, liver, kidney, and spleen (*n* = 6) of 9-week-old animals. (**B**) Violin plots showing the median allelic ratio of all replicates per tissue (*n* = 3) for autosomal genes. An allelic ratio of 1 indicates the BL6 allele, while 0 corresponds to the CAST allele. The dotted lines represent the defined cut-off values for allele-specific expression (ASE, total reads ≥ 20, allelic ratio ≥ 0.7 or ≤ 0.3). Boundaries of boxplots indicate the interquartile range around the median. Whiskers range from maximum to minimum values. Dots show gene loci with ASE. (**C**) Bar chart illustrating the proportions of allele-specific genes per tissue categorized by protein-coding status. Light grey denotes the percentage of protein-coding genes, while dark grey represents non-coding loci. (**D**) Pie chart showing biotype proportions for the allele-specifically expressed non-coding genes with available biotype information (*n* = 520, 52.74%). TEC: to be experimentally confirmed; misc RNA: miscellaneous RNAs that cannot be classified. (**E**) Correlation plot displaying the fraction of allele-specific protein-coding and non-coding genes normalized by the total number of protein-coding and non-coding loci respectively. The color code resembles the different tissue samples (*n* = 18). The correlation was calculated using a Spearman correlation (*R* = 0.66, *p* = 0.004). (**F**) Box plot illustrating the percentage of allele-specific ncRNAs nearby allele-specific or biallelic protein-coding genes (±100kb distance). Boundaries of the box indicate the interquartile range around the median. Whiskers range from maximum to minimum values. A Wilcoxon test was used to compare the allele-specific ncRNA enrichment around allele-specific and biallelic genes (*p* = 0.002). Pc: protein-coding. (**G**) Overview of the allele-specific strategy for predicting target genes of ncRNA loci based on allele-specific correlation. ncRNAs were classified as repressive or enhancing depending on their expression bias relative to nearby targets. (**H**) Schematic overview of the Allelome.LINK pipeline. The Allelome.LINK workflow starts by using Allelome.PRO2 to classify sequencing data into biallelic, maternal, or paternal expression based on reads overlapping heterozygous SNPs. Then, Allelome.LINK connects allele-specific loci within a user-defined genomic window. The output comprises a tabular list of candidate associations ranked by a linkage score and a BEDPE file to facilitate genome browser visualization. (**I**) Output of the Allelome.LINK pipeline for the lncRNA *Xist*. We used publicly available RNA-sequencing data from the placenta of E12.5 F1 hybrids (CASTxFVB *n* = 2, FVBxCAST *n* = 2,^24^). Red lines represent repressive interactions with a protein-coding gene (total reads ≥ 20, allelic ratio > 0.75 and < 0.25, window size: full chromosome). Below the X-chromosome, red lines indicate maternally expressed, while green lines show biallelic genes, including the known escape genes *Kdm6a, Eif2sx3x, Jpx*, and *Kdm5c*. (**J**) Allelome.LINK output of the *Igf2r*/*Airn* locus from placentas isolated from E12.5 embryos (*n* = 6,^12^). The upper browser track shows the repressive targets of *Airn* and their allelic ratio as depicted by the color code in the wildtype (*n* = 3, median). The arcs represent repressive interactions between *Airn* and target genes while the height resembles the linkage score. The lower panel illustrates the results for the *Airn* promoter deletion (*n* = 3, median, total reads ≥ 10, allelic ratio ≥ 0.7 or ≤ 0.3, window size: 4000kb). (**K**) Precision-recall plot evaluating the Allelome.LINK pipeline for *Airn* and the *Igf2r* locus. Precision and recall were calculated for individual replicates and combined.

### The Allelome.LINK strategy correctly assigns targets and mechanisms of imprinted lncRNAs

To facilitate the prediction of ncRNA-targets and their mode-of-action, we adapted the Allelome.PRO pipeline^19^ to make it more user-friendly and usable to multiple applications including the identification of allele-specific genomic features from individual samples or from a single cell (see Materials and methods). As an extension to Allelome.PRO2, we developed the Allelome.LINK pipeline (Figure S3). This was designed to use the allele-specific information to identify allele-specific loci within user-defined windows and predict enhancing or suppressing *cis*-effects based on allelic bias towards identical or opposite alleles, respectively (Figure 1H). We first tested our pipeline by predicting the regulatory targets of the extensively studied lncRNA *Xist*, which epigenetically silences specifically the paternal X-chromosome in extraembryonic lineages^20,21^. Using this system, we aimed to predict the targets in the placenta and accurately identified *Xist* as a repressive lncRNA for most informative X-linked genes (Figure 1I, Table S3). Notably, the pipeline did not assign repressive associations to the well-studied escape genes *Kdm6a, Eif2s3x, Jpx*, and *Kdm5c*, which are known to overcome *Xist*-mediated silencing, highlighting the robustness of our approach. We performed further validation by testing the predictions for known targets of the paternally-expressed lncRNA *Airn*, which silences genes within the *Igf2r*/*Airn* cluster in a *cis*-dependent manner^22–25^. Using wild-type and *Airn* knockout RNA-seq datasets from F1 placentas^12,26^, we identified maternal expression for known *Airn* targets and confirmed biallelic expression in *Airn* knockouts (Figure 1J). Importantly, Allelome.LINK correctly assigned *Airn* as a repressive ncRNA in the wild-type, with no linkages detected in the knockout model (Figure 1J, Table S4). Precision and recall were calculated separately for each replicate and for pooled samples. The highest precision (85.7%) was obtained by merging replicates, while the recall remained above 85% (Figure 1K). Taken together, these showcases highlight the effectiveness of the Allelome.LINK approach in predicting the *cis*-targets and mode-of-action of regulatory ncRNAs.

### Allele-specific genomics assigns targets and mechanisms to a significant fraction of ncRNAs across mouse organs

After validation, we applied the Allelome.LINK approach to the generated allele-specific mouse bodymap (Figure 2A). On average, we predicted 66.2 linkages per tissue, with the spleen showing the highest number of ncRNA-mRNA ASE events (*n* = 99). Notably, there was a higher number of predicted enhancing than repressive linkages across all organs (Figure 2B). The well-established imprinted interaction between *Airn* and *Igf2r* was one of the top linkages in all tissues except the brain, confirming previous findings and further validating our strategy^12^ (Figure 2C). In total, the Allelome.LINK approach successfully linked an average of 11.3% of the allelic ncRNAs to their putative target genes (Figure 2D). Notably, the majority of linkages were tissue-specific (62.2%, *n* = 247), while only 37.8% (*n* = 150) were observed in two or more tissues (Figure 2E). Interestingly, while repressive linkages showed an evenly distributed target distance, with the highest peak observed around 32kb, enhancing links control targets in close proximity, suggesting regulation from bidirectional promoters (Figure 2F and 2G). We selected two examples of high-confident linkages to highlight the target gene prediction based on the allelic read counts. First, a repressive anti-sense link between the ncRNA *Gm35993* and the pcGene *Acmsd* detected in the kidney (Figure 2H, left panel), and second, an intergenic repressive link identified in the liver, where *Gm38596* is predicted to suppress *Sult2a7* (Figure 2H, right panel). In summary, our analysis yielded a total of 397 predicted ncRNA-mRNA ASE events across all tissues using stringent criteria (Figure 3, Table S5). To facilitate further exploration and validation of the predicted associations, we have made all candidate links available for visual inspection, providing a valuable resource for the research community (see Data availability).

**Figure 2.**
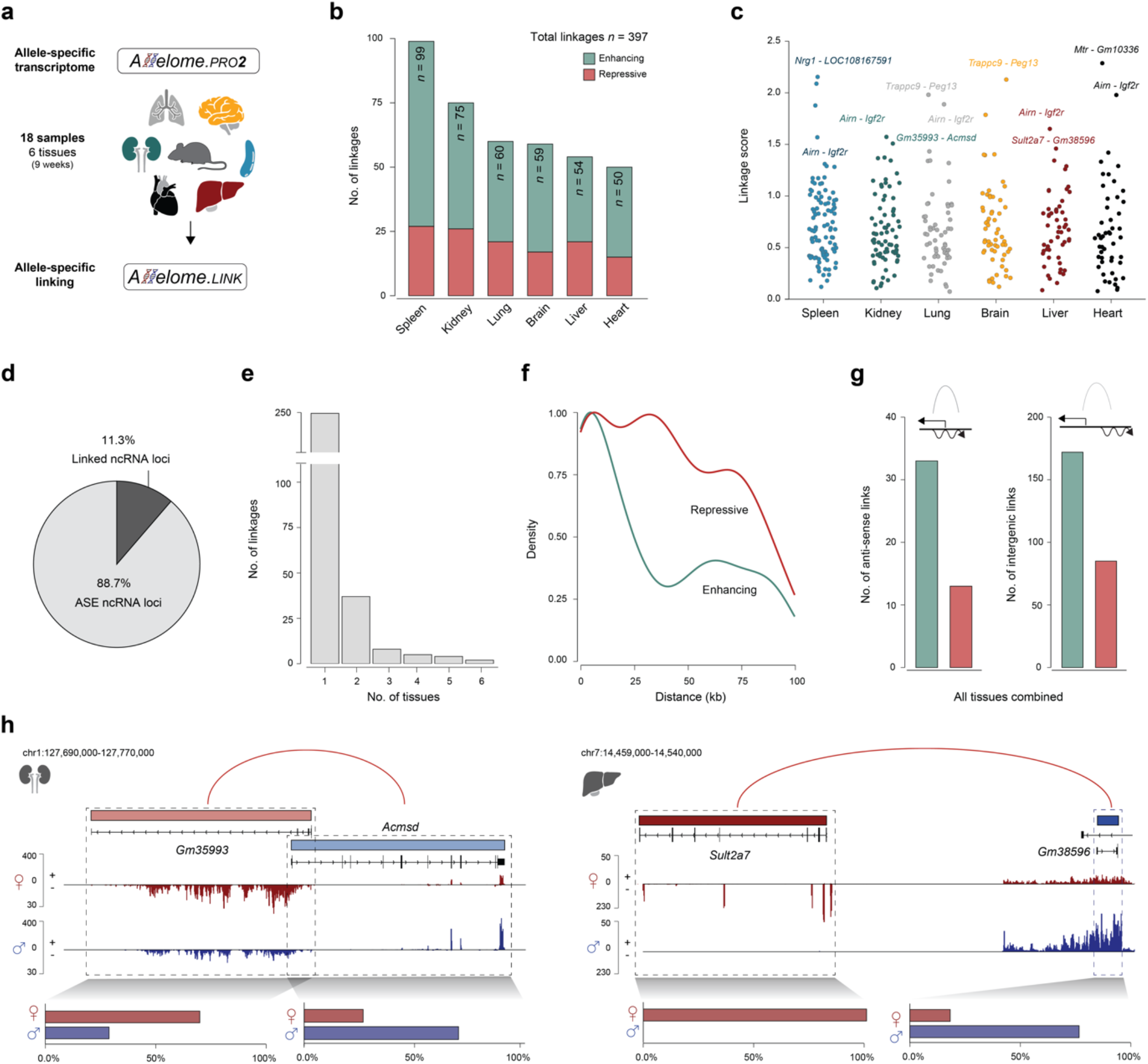
Allelome.LINK elucidates shared and tissue-specific ncRNA to protein-coding target links and their mode-of-action. (**A**) Schematic overview of the Allelome.LINK workflow in mice. Candidate linkages were predicted using the allele-specific transcriptome data of the brain, heart, lung, liver, kidney, and spleen (*n* = 6). (**B**) Number of linkages identified per tissue. Green denotes the enhancing interactions, while red represents repressive linkages. The numbers display the total abundance of linkages per tissue. (**C**) Manhattan plot showing the distribution of linkages per tissue along the linkage score. (**D**) Pie chart displaying the mean proportion of allele-specific ncRNA loci per tissue. On average, 11.3% of the informative ncRNAs with an allelic bias could be linked to their putative targets. (**E**) The number of shared linkages across the six different tissues. (**F**) Density plot, illustrating the distribution of distances between ncRNAs and assigned target genes. The density curves are separated for enhancing (green) and repressive (red) interactions. The dashed lines represent the median distance for enhancing and repressive links, respectively. (**G**) Bar plots showing the number of enhancing (green) and repressive (red) links across all samples. The left panel depicts anti-sense, while the right panel shows non-overlapping associations. (**H**) Examples of two high-confident linkages displayed in the IGV browser. The upper part shows the Allelome.LINK output, with red arcs representing repressive linkages. Gene colors indicate the allele-specific bias towards maternal (red) and paternal (blue) expression. Below, the panel shows strand- and allele-specific mapping of sequencing reads for the maternal and paternal allele. Bar plots quantify mapped reads for each allele. Left: Repressive anti-sense linkage between *Gm35993* and *Acmsd*, detected in the kidney. Right: Intergenic linkage identified in the liver, where *Gm38596* is predicted to suppress *Sult2a7*.

**Figure 3.**
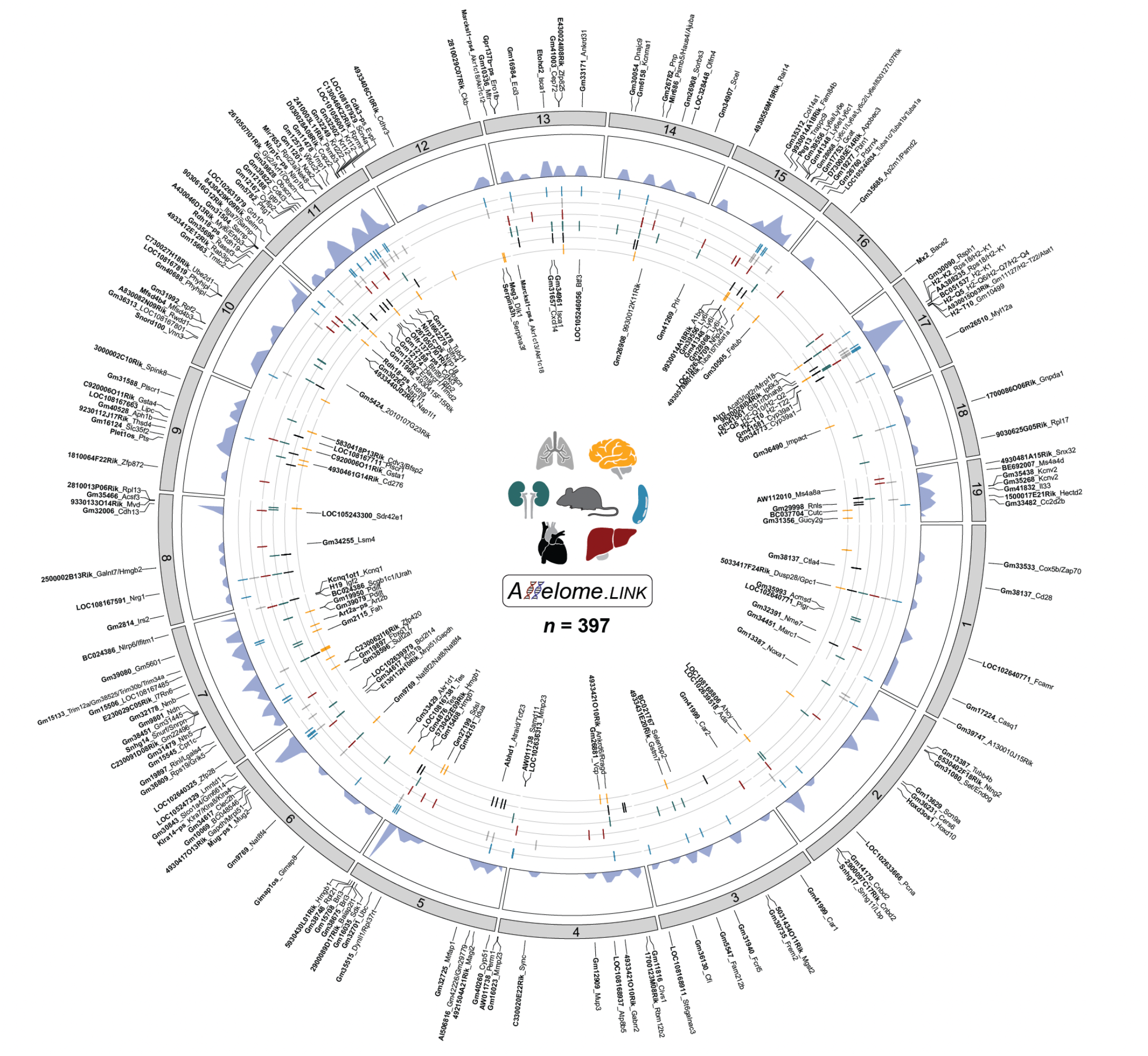
Comprehensive mapping of 397 identified candidate interactions between ncRNAs and protein-coding genes in mice. Candidate interactions were visually represented in a chord plot spanning chromosome 1-19. Linkages on the outside indicate enhancing interactions, while internal linkages resemble repressive links. For each linkage, the ncRNA is highlighted in bold font and the target is separated by an underscore. For ncRNAs with multiple targets, the protein-coding genes are separated by dashes. Density plots show the distribution of linkages across chromosomes. The genomic location of the ncRNA is shown in the inner circle, with each line corresponding to a tissue (spleen: blue, lung: grey, liver: red, kidney: green, heart: black, brain: yellow).

### Exploiting inter-individual variation in humans elucidates targets and mechanisms of ncRNAs across tissues

To extend the approach beyond mouse models, we applied our linking strategy to allele-specific haplotype data of the Genotype-Tissue Expression resource (GTEx)^17^. This comprehensive dataset includes 153 million haplotype measurements across 15,253 samples and 54 human tissues of nearly 1,000 individuals (Figure 4A). As the GTEx data is not strand-specific, we removed gene loci with overlapping exons to mitigate the allelic bias of overlapping transcripts. This process yielded 1825 informative ncRNA and 6281 pcGene loci, comprising 924,440 and 27,155,698 allele-specific measurements, respectively (*n* = 8106 loci, total read cut-off ≥20, Figure S4). On average, we detected 3,580 (ncRNAs *n* = 312, pcGenes *n* = 3,268) unique loci with ASE per sampling site. Generally, we found that the number of allele-specific loci increased with the number of individuals (Figure 4B, Table S6). For instance, the kidney-medulla showed the lowest number of ASE genes (*n* = 310; 4 individuals), while the lung showed the highest number (*n* = 4,983; 515 individuals). In total, we obtained informative haplotype data for 1,825 unique ncRNAs across all sampling sites.

**Figure 4.**
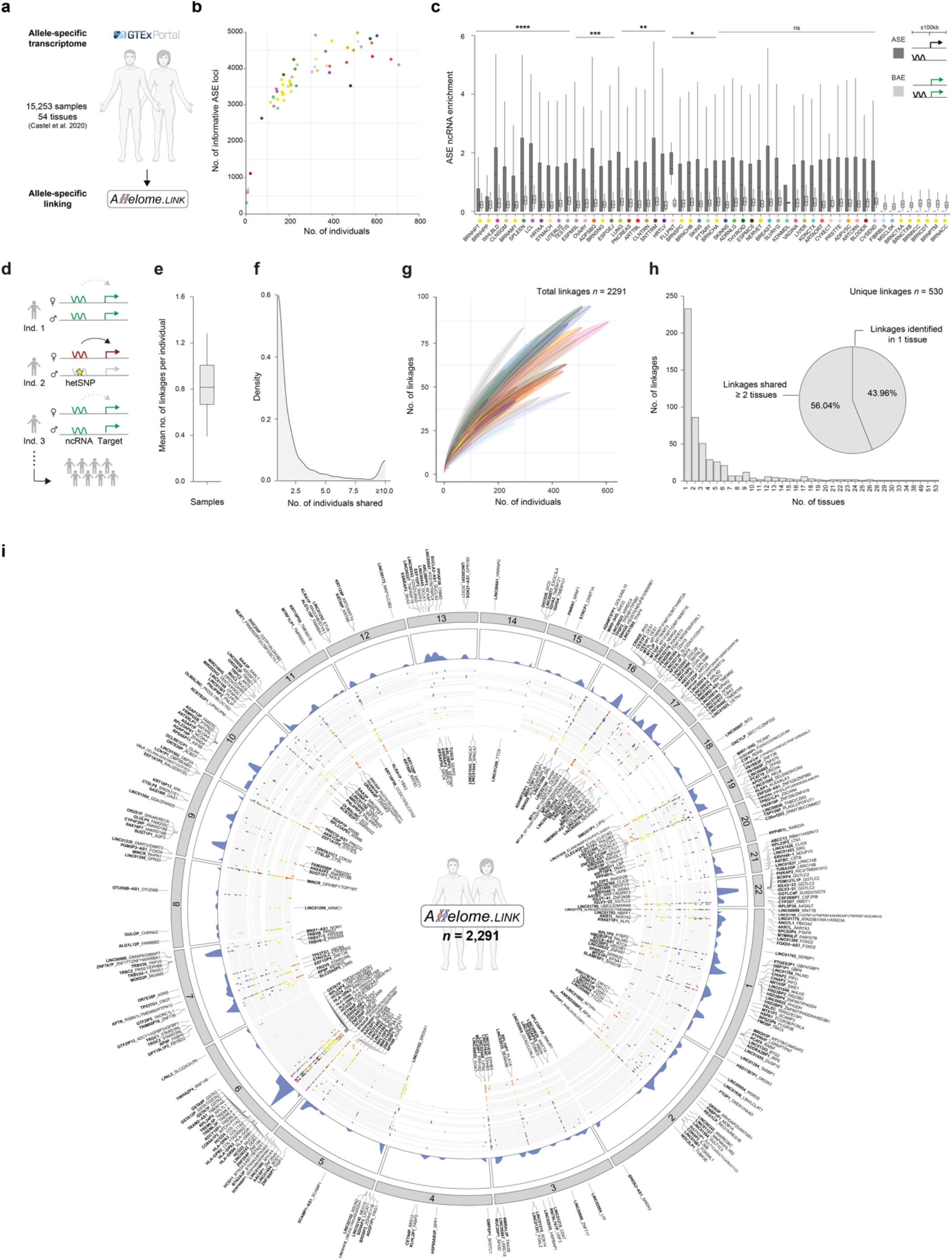
Applying the Allelome.LINK approach to the GTEx database predicts 2,291 ncRNA-mRNA ASE events in humans. (**A**) Schematic overview of predicting human ncRNA-mRNA ASE events using the GTEx database. Allelome.LINK was applied to the publicly available data generated by Castel et al. (GTEx v8 release). This dataset contains 153 million allele-specific haplotype measurements across 15,253 samples and 54 different human tissues^17^. (**B**) Scatter plot showing the number of individuals along the number of unique allele-specific loci after removing ncRNAs ≤ 200bp in exon length and overlapping genes. Informative loci (total read number ≥ 20) were considered allele-specific with allelic bias ≥ 0.7 or ≤ 0.3 (n = 193,327). The color code represents the different sampling sites. (**C**) Enrichment of allele-specific ncRNAs around protein-coding genes. Box plots show the proportions of allele-specific ncRNAs around allele-specific (dark grey) and biallelic (light grey) protein-coding genes within ±100kb distance, per individual and tissue. Boundaries of the box indicate the interquartile range around the median. Whiskers range from maximum to minimum values. Wilcoxon tests were used to compare allele-specific ncRNA enrichment around allele-specific and biallelic genes. The number of asterisks indicates the significance level. ASE: allele-specific expression, BAE: biallelic expression. (**D**) Schematic overview demonstrating how genetic diversity in humans leads to the discovery of novel linkages. While the enhancing interaction is evident in all three individuals, only individual 2 allows the detection of the association due to a heterozygous SNP that induces allele-specific expression of the ncRNA. In individuals 1 and 3, this interaction is masked by the biallelic expression of the ncRNA. (**E**) Mean number of linkages per individual across all tissues. Boundaries of the box indicate the interquartile range around the median. Whiskers range from maximum to minimum values. (**F**) Density plot illustrating the number of individuals that share a linkage. (**G**) Line plot showing the mean number of linkages, along with the number of individuals. Means and standard deviations were calculated by random sampling with 1000 iterations. The color code corresponds to the different sampling sites. (**H**) Bar plot illustrating the fraction of linkages that are shared across a diverse number of tissues. The pie plot resembles the proportions of the 530 unique linkages that are tissue-specific and shared in ≥ 2 tissues. (**I**) Chord plot of human candidate interactions spanning chromosomes 1-22. Linkages on the outside of the chord plot indicate interactions that were detected as enhancing across most samples, while internal linkages resemble likely repressive links. For each interaction, the ncRNA is highlighted in bold font and the target is separated by an underscore. Density plots show the distribution of linkages across chromosomes. The genomic location of the ncRNA is shown in the inner circle, with each line corresponding to a tissue according to the color code.

To validate the non-random association between allele-specific ncRNAs and proximal allelic pcGenes in humans, we quantified their co-occurrence within a distance of ±100kb per individual and tissue (Figure 4C). In line with findings in mice, we observed a significant enrichment of allele-specific ncRNAs in the proximity of allelic pcGenes for half of the investigated tissues (27 of 54, Figure 4C, Table S7). Hence, we proceeded to leverage the Allelome.LINK approach on the extensive allele-specific dataset, allowing us to predict human ncRNA-mRNA ASE events. Given the outbred nature of humans, each individual is expected to have a unique allele-specific landscape shaped by their personalized set of genetic variants. This diversity provides an opportunity for the discovery of novel ncRNA-mRNA ASE events, with each individual potentially revealing further linkages. Notably, the ncRNA mechanisms and targets identified through these individual-specific linkages are expected to be common across individuals but can only be detected in individuals with a genotype that leads to allelic bias (Figure 4D). On average, each individual revealed approximately one linkage per sampling site (Figure 4E). The majority of the linkages (63.77%) were unique to a single individual, while 36.23% were detected in multiple samples (Figure 4F, Table S8). To test whether each individual contributes to new linkages and whether we already reached saturation in this large dataset, we generated a saturation curve for each tissue. Intriguingly, we observed that each sample contributes to identifying novel links, as we did not observe a saturation at any sampling site (Figure 4G, Table S9). This finding highlights the power of this approach and the future potential of adding more individuals to uncover new associations by exploiting the genetic variation inherent in the human population. Consistent with the findings in mice, our analysis revealed many tissue-specific linkages, accounting for 43.96% (*n =* 233), while 56.04% (*n* = 297) were shared across two or more tissues (Figure 4H, Table S10). These findings underscore the extensive tissue-specific regulatory roles of ncRNAs. Overall, we identified 2,291 human ncRNA-mRNA ASE events across all tissues, comprising 324 unique linked ncRNAs (Figure 4I, Table S11). Hence, we predicted the *cis*-targets of 17.75% of the informative ncRNA loci using the allele-specific approach. This resource provides insight into the complex interplay between human ncRNAs and their targets and serves as an ideal starting point to select candidates based on target function (see Data availability).

### The majority of identified human linkages are confirmed by eQTL data

In order to confirm the predicted ncRNA-target gene associations, we overlapped the linked ncRNA loci with eQTL data that derived from the same samples, containing 21,412,255 fine-mapped eQTLs of 49 tissues^27^. eQTL data was absent for the Kidney - Medulla, Fallopian Tube, Cervix - Endocervix, Cervix - Ectocervix, and Bladder. Remarkably, we confirmed an average of 77.47% (sd = 9.83) of the linkages with eQTLs across all 49 sampling sites (Figure 5A). Of these eQTLs, a mean of 18.72% of our ncRNA-mRNA ASE events were confirmed in the corresponding tissue, with tibial nerves showing the highest eQTL support of 35.7% (Table S11). To validate our regulatory mechanism predictions, we assessed the proportion of enhancing and repressive assignments for linkages shared across individuals. Initially, we tested the mechanism prediction for genes in the HLA cluster, which are known for high genetic variation and different allelic statuses among individuals^28^. As expected, we observed a random distribution of mechanisms across individuals with 49.25% enhancing and 50.75% repressive linkages (Figure 5B). Additionally, we tested our strategy on the imprinted *MEG3*/*DLK1* loci, where the lncRNA *MEG3* is known for maternal-mediated *DLK1* repression^29^. This epigenetic interaction provides a reliable model for assessing consistency across individuals, as the repressive interaction is independent of individual genotypes. We analyzed tissues where imprinting of *MEG3* and *DLK1* has been previously confirmed^16^ and correctly identified 79.46% of 564 linkages as repressive (Figure 5C). In conclusion, the analysis validated a substantial proportion of the predicted linkages by eQTL data and demonstrated the reliability of our approach to accurately assign mechanisms to distinct ncRNA loci across individuals.

**Figure 5.**
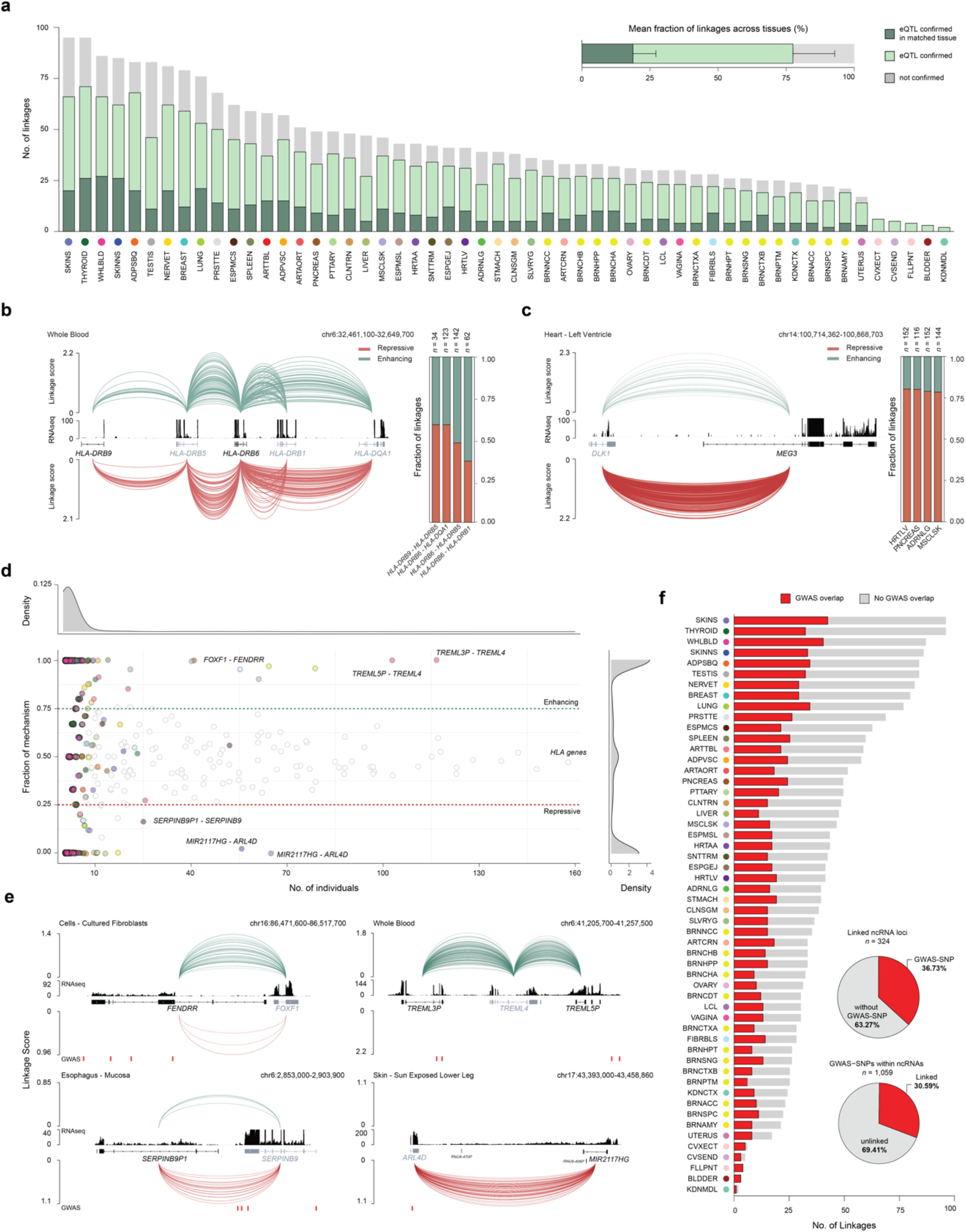
Validating the anticipated ncRNA-mRNA ASE events and linking GWAS SNPs in the non-coding genome to their protein-coding targets. (A)Bar chart illustrating the percentage of eQTL validation for the identified linkages per tissue. Linkages were classified as confirmed if an eQTL that overlaps the ncRNA gene body affects the expression of the linked target gene. Dark green represents the percentage of linkages confirmed by an eQTL within the same tissue, while light green represents the percentage of linkages validated by eQTLs identified in different tissues. The fraction of linkages that could not be confirmed by eQTLs is depicted in grey. Bar plot above showing the mean fraction of eQTL validated linkages across all tissues (*n* = 49). Error bars indicate the standard deviation. (**B**) Allelome.LINK output for the HLA genes: *HLA-DRB9, HLA-DRB5, HLA-DRB6, HLA-DRB1, HLA-DQA1* in whole blood samples (*n* = 611). Green arcs resemble enhancing (*n* = 178), while red arcs indicate repressive linkages (*n* = 183). The height of the arcs is proportional to the linkage score. The RNA-sequencing track was obtained from GTEx and represents a single sample of the tissue. Gene names are colored according to the coding status (black: non-coding, grey: coding). The bar plot illustrates the fraction of mechanism (green: enhancing, red: repressive) for each of the four linkages across individuals. The *n* number depicts the total number of individuals where the link was detected. (**C**) Allelome.LINK output as in (b) for *MEG3* and *DLK1* across Heart - Left Ventricle samples (*n* = 386). The bar plot illustrates the fraction of mechanism (green: enhancing, red: repressive) for the *DLK1* - *MEG3* interaction in the tissues: Heart - Left Ventricle, Pancreas, Adrenal gland, and Muscle - Skeletal. (**D**) Scatter plot showing the fraction of enhancing or repressive mechanisms along the number of individuals. The mechanism fraction is calculated by determining the number of individuals that show a specific linkage as enhancing, to assess consistency across samples within a tissue. 1 indicates all individuals show the linkage as enhancing, while 0 means all detections were repressive. The color code is according to the tissue. Dots without color represent linkages that include genes of the HLA cluster. Density plots summarize linkages across tissues, illustrating their distribution based on the number of individuals and the proportion of mechanisms observed. (**E**) IGV-browser examples showing the Allelome.LINK output as in (b) and (c), showcasing high-confident examples detected in (d). Three enhancing (Cells - Cultured fibroblasts: *FENDRR* - *FOXF1*, Whole Blood: *TREML5P* - *TREML4, TREML3P* - *TREML4*) and two repressive linkages (Esophagus - Mucosa: *SERPINB9P1* - *SERPINB9*, Skin - Sun- Exposed Lower Leg: *MIR2117HG* - *ARL4D*) are highlighted. Below, red lines mark GWAS hits. (**F**) Bar plot illustrating the number of linkages, where the ncRNA overlaps with a GWAS-SNP per tissue. The upper pie chart illustrates the proportion of linked ncRNA loci that overlap GWAS-SNPs out of the total number of linked ncRNAs (*n* = 324). The lower pie chart shows the total number of GWAS-SNPs that overlap informative ncRNAs (*n* = 1059), highlighting the fraction of SNPs that could be linked to a protein-coding target (*n* = 324).

### Discovery of high-confidence ncRNA linkages through over-representation of targets and mechanisms across individuals

To elucidate high-confidence ncRNA-target linkages, we took advantage of the large number of individuals to determine how many links are represented in many individuals that consistently show repressive or enhancing mechanisms. Applying a consistency cut-off of ≥ 75%, requiring a linkage to be detected as either enhancing or repressive in at least 75% for a given tissue, we identified 35.0% repressive (*n* = 802) and 48.1% enhancing (*n* = 1,102) linkages (Figure 5D, Table S11 and S12). The mechanisms for the remaining predictions were randomly assigned (16.89%), with a significant proportion (36.69%) encompassing the HLA genes (Figure 5D). Next, we investigated all linkages present in more than 10 individuals for a given tissue and identified 24 high-confident predictions (repressive *n* = 9, enhancing *n* = 15) that showed consistent mechanism prediction (≥ 75% enhancing or repressive). Among those linkages, we highlight the interaction between *FOXF1* and the lncRNA *FENDRR*, detected in 55 cultured fibroblast samples, with 52 consistently showing enhancing effects (Figure 5D and 5E). This finding in humans is supported by a previous study in mice that identified a feedback loop between *Fendrr* and *Foxf1*^30,31^. Another example is the pcGene *TREML4*, which is predicted to be positively associated with the two ncRNAs, *TREML3P* and *TREML5P*, supported by 117 and 103 enhancing interactions in whole blood, respectively (Figure 5D and 5E). Additionally, we note the repressive linkage between the ncRNA *SERPINB9P1* and the target gene *SERPINB9 (n =* 21), as well as the ncRNA *MIR2117HG* and the pcGene *ARL4D (n =* 65, Figure 5D and 5E). Due to the high level of agreement of the predicted mechanism across multiple individuals, these linkages anticipate a high likelihood of true regulatory associations. This highlights the potential of our strategy to enrich for functional ncRNAs as more individuals are sequenced.

### GWAS integration permits assignment of non-coding risk variants to pcGene targets

Interestingly, we noticed that our high-confident interactions overlapped with GWAS variants in either the ncRNA or the pcGene locus (Figure 5E). Therefore, we used all GWAS variants documented in the NHGRI-EBI catalog to elucidate the functional role of non-coding risk variants^32^. We filtered for variants overlapping our informative ncRNA loci maintaining 1,059 variants, which we intersected with the linked ncRNAs. Notably, a significant fraction of the linked ncRNAs harbored at least one GWAS variant across different tissues (36.73%, *n* = 119 ncRNAs, Figure 5F). Skin samples showed the highest number of linkages, encompassing a risk variant within a ncRNA locus (*n* = 42), followed by samples of the whole blood (*n* = 40, Figure 5F). Overall, our approach assigned 30.59% (*n* = 324) of the ncRNA-overlapping GWAS variants to their protein-coding targets (Figure 5F). All ncRNA-mRNA ASE events and the integrated GWAS overlap have been made publicly available to enable further exploration (see Data availability, Table S11). These results provide valuable insights into the disease-related landscape of regulatory ncRNAs and associated risk variants in the relevant tissues. With a growing pool of GWAS data and individuals being sequenced, our strategy has great potential to identify the functional targets of risk variants located in the non-coding genome.

## Discussion

A significant proportion of disease variants are located in ncRNA loci, which serve as important regulators of the genome^8,33^. Thus, linking ncRNA loci to their targets will provide critical insights for a better understanding of complex diseases. Connecting ncRNAs to their target genes presents significant challenges, primarily due to their dynamic functionality across temporal and spatial variations. This includes diverse developmental stages, tissues and cell types, and responses to environmental cues^10,34^. To address this challenge, we used ASE, which avoids the dynamic nature by comparing allelic expression levels within identical environments. To date, numerous studies in a wide range of species, including humans, mice, plants and yeast, have used ASE to identify the contribution of regulatory variants to gene expression^35^. However, instead of focusing on the individual effects of variants, our strategy identifies co-occurring ASE loci, providing insights beyond variant-specific effects. Given the rarity of ASE events, our discovery of significant clustering of allele-specific ncRNAs around allele-specific pcGenes, along with a high validation rate, underscores the genuine co-regulatory interplay between allele-specific ncRNAs and neighboring allelic pcGenes. Using this concept, our study identified 397 mice and 2,291 human ncRNA-mRNA ASE events and their regulatory mode-of-action. This extensive resource in mice and humans serves as an ideal starting point for the community to select candidates for further investigation and validation.

Given that our approach involves comparing alleles within the same cellular environment, the predicted gene regulation of linkages is limited to *cis*-effects. This approach allowed us to assign whether a ncRNA has a repressive or activating role for overlapping, antisense or long-range targets. While our transcriptomic data points to an RNA mediated mechanism, we cannot exclude a role for DNA regulatory elements within the ncRNA locus, or the act of transcription, both capable of modulating the expression of adjacent genes independently of the transcribed RNA^33,36,37^. Moreover, various regulatory mechanisms in ncRNA loci can simultaneously impact the expression of a target gene, including both *cis*- and *trans*-regulatory effects^36,38^. A recent study by Tsouris et al. (2024) proposed comparative analyses of multiple parent-hybrid trios to evaluate the relative impact of *cis*-versus *trans*-effects on a given locus^39^. Integrating this approach with the identified linkages could further delineate the *cis*-effects of our linked ncRNAs. In addition, the integration of diverse mouse strains provides the opportunity to uncover novel associations and cross-validate existing predictions. This concept can also be translated to various conditions, such as aging and disease, offering additional insights into the biology of ASE and potential regulatory shifts.

Expanding our scope to humans, our results showed that the majority of linkages were identified in single individuals, consistent with the reported assumption that most ASE is driven by genetic variation^13,15^. We evaluated the mechanism assignment on the imprinted *MEG3*/*DLK1* loci and observed a correct prediction of 79.50%. The remaining misassignment rate is likely due to phasing inaccuracies that increase with distance. The outbred nature of the human population results in a distinctive allele-specific landscape for each individual, shaped by their unique set of genetic variants. This inherent diversity provides the opportunity to uncover novel associations between ncRNAs and their targets with each individual. It is important to note that the ncRNA mechanisms and targets identified by these individual-specific associations are expected to be common across individuals, but can only be detected in individuals with a genotype that leads to allelic bias. In addition, we identified a subset of linkages that we defined as high-confidence examples based on their ability to replicate across individuals with common mechanisms-of-action. More individual sequencing data is expected to reveal new associations and help to identify highly probable associations with common mechanisms.

Finally, by integrating GWAS data, we were able to link 30.59% of the variants that overlap linked ncRNA to their pcGene. The continued accumulation of individual sequencing data and non-coding risk variants provides great potential to increase this fraction in the future. In our study, the absence of strand-specific RNA-seq data in humans required the removal of overlapping transcripts to avoid false positives, reducing the number of informative ncRNAs. It is also important to note that another large proportion of non-coding GWAS variants are located in regulatory DNA elements, such as enhancers or repressors^2^. Nevertheless, our strategy can also be used to link DNA regulatory elements and their associated disease variants to their nearby targets, for example by providing ATAC-sequencing as input. In conclusion, our approach provides a novel framework to uncover the targets and mode-of-action of the non-coding genome. With the future availability of more extensive individual sequencing data and non-coding risk variants, our strategy will illuminate the vast landscape of functional *cis*-acting elements, thereby decoding the intricate complexity of the non-coding genome and its role in complex diseases.

## Acknowledgements

We thank John Rinn and Life Science Editors for comments on the manuscript. Sequencing was conducted at the Helmholtz Munich Genomics Core Facility. This work was supported by the Deutsche Forschungsgemeinschaft 389 (Project-ID: 403584255—TRR 267) and a DZHK (Deutsches Zentrum für Herz-Kreislauf Forschung) Junior Research Group grant. This paper was typeset with the bioRxiv word template by @Chrelli: www.github.com/chrelli/bioRxiv-word-template

## Author contributions

Tim Hasenbein: Conceptualization, Data curation, Software, Formal analysis, Investigation, Visualization, Methodology, Writing original draft, Writing—review and editing. Sarah Hoelzl: Investigation, Resources, Methodology. Stefan Engelhardt: Writing—review and editing. Daniel Andergassen: Conceptualization, Supervision, Funding acquisition, Project administration, Data curation, Investigation, Formal analysis, Validation, Methodology, Writing—original draft, Writing—review and editing.

## Data availability

All ncRNA-mRNA ASE events for mice and human are provided in the Supplementary Table. Further, we provide access to all linkages via the Integrative Genomics Viewer to present the target and mechanism predictions for each ncRNA for the community to explore. This resource can be accessed at https://github.com/AndergassenLab/Allelome.LINK/tree/main/02_resource. The source code for Allelome.PRO2 and the Allelome.LINK pipeline will be available at the same GitHub page upon publication.

## Materials and Methods

### Mouse strains

To generate F1 hybrids, BL6 females (JAX: Strain #000664) were paired with CAST males (JAX: Strain #000928). All mice were housed in an open-cage environment provided by the Technical University Munich Institute of Pharmacology and Toxicology. Animal experiments of the study were conducted according to the German Animal Welfare Act (Tierschutzgesetz and Tierschutzversuchstierverordnung) and EU guideline 2010/63. Approval was received from the relevant authorities (Kreisverwaltungsreferat der Stadt München, Veterinäramt München Stadt, permit granted according to §11, Paragraph 1 sentence 1 no. 1 of the German Animal Welfare Act).

### Tissue isolation and library preparation

Three replicates were harvested for brain, heart, liver, lung, kidney, and spleen from 9-week-old F1 hybrid mice (BL6 x CAST) and snap-frozen in liquid nitrogen. The GentleMACS Dissociator (program RNA_02_01) was used to homogenize tissues in 1 ml TRIzol per 50-100 mg sample. RNA isolation was conducted as described in the manufacturer’s instructions (Invitrogen, TRIzol Reagent, Cat. #15596018) using 1 ml of the sample solution. RNA-seq libraries were constructed with 100 ng of RNA and Illumina’s Stranded mRNA Prep Ligation Kit. The Agilent TapeStation System was used to assess the concentration and fragment length of the libraries. Sequencing was performed using a NovaSeq6000 50bp paired-end sequencing.

### Preprocessing of RNA-seq data

Paired-end RNA-seq data of 9-week-old mice were aligned to the GENCODE_M25GRCm38.p6_201911 primary assembly using STAR (v2.6.0c,^40^). Reads characterized by intron sizes exceeding 100000, multimappers, and alignments containing non-canonical junctions were removed prior to subsequent analysis. Uniquely aligned reads were quantified using htseq-count and the *–stranded reverse* flag (HTSeq version 0.11.3,^41^). Transcripts per million (TPMs) were calculated by custom R scripts for sample clustering based on Pearson correlation. Aligned bam files were further split into forward and reverse strands using a custom Perl script. BigWig files were generated using bam2wig.py.

Public placental E12.5 RNA-seq data (2x CASTxFVB, 2x FVBxCAST) was downloaded from the GEO database for SRR3085950, SRR3085951, SRR3085952, SRR3085953^12^, as well as for the *Airn* knockout (SRR8753471, SRR8753472, SRR8753473, SRR8753474 SRR8753475, SRR8753476^24^). The alignment was performed as described above, except for the *Airn* deletion data, which was aligned in single-end mode.

### Allocation of the sequencing reads to the alleles using SNPsplit

Sequencing reads were assigned to the parental alleles using SNPsplit v0.3.2^42^. Fastq files were merged across replicates per tissue to increase gene coverage. Subsequently, reads were mapped to an N-masked genome generated with *SNPsplit_genome_preparation* and STAR (v2.6.0c) for BL6/CAST SNPs, which were obtained from the Sanger database (mgp.v5.merged.snps_all.dbSNP142.vcf,^43^). The same annotation file was used for alignment, but using the settings: *–alignIntronMax 100000, – outFilterIntronMotifs RemoveNoncanonical, –outFilterMultimapNmax 1, – alignEndsType EndToEnd, –outSAMattributes NH HI NM MD*. The resulting aligned bam file was separated by strand as described and used as input for SNPsplit v0.3.2.

### Generation of the Allelome.PRO2 pipeline

We developed Allelome.PRO2, an updated version of the previously published pipeline^19^ to facilitate the usage of single samples. Unlike its predecessor, Allelome.PRO2 does not discriminate between allele-specific loci arising from imprinted or genetic factors. This update expands the identification of ASE at the individual level, improving its applicability to diverse biological samples, including human datasets where forward and reverse crosses cannot be obtained. The source code for Allelome.PRO2 will be publicly available at https://github.com/AndergassenLab/Allelome.LINK/ upon publication. We updated the usage of the pipeline by simplifying the configuration process. The config file has been removed and substituted by command-line parsing via input flags, allowing the user direct execution without prior configuration. Additionally, the following input requirements have been removed to implement the one-sample approach and facilitate the usage of the pipeline: *ratio, fdr_param, main_title, y_axis, strains, for_c1, for_c2, rev_c1, rev_c2*. To improve computational time, a new input flag was introduced to directly specify a total read cut-off, which removes loci with insufficient SNP coverage. Furthermore, we revised the classification scheme for Allelome.PRO2 categories such as *Imprinted: Maternal (MAT), Imprinted: Paternal (PAT), Strain bias: Strain 1, Strain bias: Strain 2, Not informative (NI), and No SNP (NS)* have been removed. This update allows users to obtain the allele-specific information of individual samples and makes the requirement of four samples redundant. Along with these changes, we also removed the FDR-based mock comparison and the user-set ratio filter needed for classification. As a result, Allelome.PRO2 provides allelic ratios for each locus without classification, allowing the pipeline to be run on individual samples. This refinement has led to the removal of several output files, including *<name>_IG*.*txt, <name>_SG*.*txt, <name>_locus_full*.*txt, <name>_SNP_full*.*txt, <name>*.*pdf*, and *info*.*txt*. A comprehensive log file has been implemented for better tracking and debugging. Additionally, we integrated *pileup_filter*.*pl, read_count*.*pl, bed_creator_SNP*.*sh*, and *bed_creator*.*sh* in the score.R script to streamline the pipeline’s functionality.

### Allele-specific analysis of mice samples

Allele-specific analysis for RNA-seq data was performed using the updated Allelome.PRO2 pipeline. We used the RefSeq gene annotation which was further split into forward and reverse strand. To analyze the transcriptomic bodymap of nine-week-old F1 mice we obtained 20,635,313 SNPs between C57BL_6NJ x CAST_EiJ from the Sanger database (mgp.v5.merged.snps_all.dbSNP142.vcf,^43^). Allelome.PRO2 was run individually for each strand per sample, with a minimum read cut-off ≥ 1 per SNP. Strands were merged before pooling individual replicates. We retained loci with ≥ 20 total reads per gene across all three replicates only. Reads and allelic ratios were summarized using median values, while the minimum value was selected for the allelic score to ensure robust allele-specific calling. Genes located on the X-chromosome were removed prior to subsequent analyses in order to mitigate allele-specific bias due to skewed X-inactivation.

To get the allele-specific information of X-linked genes expressed in the placenta, we used the publicly available data from ^12^ and followed the described processing steps with the exception of using a SNP file generated from mgp.v5.merged.snps_all.dbSNP142.vcf for CAST x FVB (*n* = 20,581,027). Replicates from forward and reverse crosses were pooled.

Due to the lack of strand-specific information, unstranded files were used for the *Airn* knockout data, along with previously published CAST/FVB SNP file, which included 16,988,479 variants^24^. Replicates were pooled as described, using a total read count threshold ≥ 10.

### Development of the Allelome.LINK extension tool

To streamline the prediction of ncRNA-targets and their mode-of-action, we developed Allelome.LINK as an extension of Allelome.PRO2. Utilizing allele-specific data, this tool connects ASE loci in *cis* within user-defined genomic windows. As input, Allelome.LINK requires a text file formatted as the locus_table.txt output of Allelome.PRO2. The tool then filters loci with ASE values exceeding the specified cut-off threshold. Biased loci are intersected and linked if they co-occur within the defined genomic window. Mode-of-action is determined as enhancing or repressive based on allelic correlation or anti-correlation. A score is calculated for each linkage using the formula:

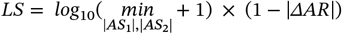

Where AS is the allelic score for each locus as calculated by Allelome.PRO and ΔAR the differences in the allelic ratio calculated as

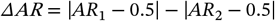

This definition is based on the assumption that regulatory genes correlate in their allelic bias. Each allelic ratio is standardized by subtracting 0.5 to center the values. Consequently, a small ΔAR signifies closely matched allelic ratios, irrespective of whether the bias is toward identical or opposite alleles. The allelic score is further implemented in the scoring calculation, assigning higher linkage scores to interactions characterized by robust allelic biases. This scoring accounts for the statistical significance derived from the *p*-value of a binomial distribution as described in ^19^. As output, Allelome.LINK provides the linkage table ranked by linkage scores as well as BEDPE and BED files for intuitive visualization via genome browsers. Enhancing interactions are highlighted in green, while repressive predictions are marked in red. Additionally, each analysis run generates a comprehensive log file. The source code will be publicly available at https://github.com/AndergassenLab/Allelome.LINK/ upon publication.

### Data analysis using Allelome.LINK with mice samples

To link ncRNAs to their putative protein-coding targets in mice, we used the mapped allelome of adult organs (spleen, kidney, lung, heart, liver, and brain) obtained from 9-week-old F1 hybrid mice. We applied Allelome.LINK using a total read cut-off ≥ 20 and an allelic ratio cut-off ≥ 0.7 or ≤ 0.3. The genomic window was set to ±100kb due to the significant enrichment observed within this range. Subsequently, we filtered for non-coding to protein-coding interactions for further downstream analysis. For the prediction of *Xist* targets, we used an allelic bias ≥ 0.75 or ≤ 0.25 along with a chromosome-wide window. The genomic range was also adjusted for the *Airn* knockout data to ±4000kb to encompass the entire *Airn/Igf2r* cluster. Allelic bias criteria were maintained at 0.7/0.3.

### Linking ncRNA-mRNA ASE events using human samples of the GTEx database

Human allele-specific measurements were derived from publicly available data generated by ^17^ from the GTEx v8 release (phASER_WASP_GTEx_v8_matrix.gw_phased.txt). This large dataset contains 153 million allele-specific haplotype measurements across 15,253 samples from 54 human tissues. The haplotype data was merged with the sample information (GTEx_Analysis_v8_Annotations_SampleAttributesDS.txt) and formatted using R to match the Allelome.PRO2 output. Because the GTEx data lack strand-specificity, we filtered out ncRNAs with a combined exon length ≤ 200bp and overlapping gene loci. To ensure data consistency, we used the GENCODE v26 annotation, to match the annotation of the resource. After filtering the annotation, we maintained 21,921 loci, including 14,056 ncRNAs and 7,865 pcGenes. A gene was classified informative with a total read cut-off ≥ 20 in at least one sample. Subsequently, we ran Allelome.LINK for each individual using an allelic ratio cut-off ≥ 0.7 or ≤ 0.3 and a window size of ±100kb.

### Validation of linkages using eQTL information

To validate our predictions of ncRNA interactions with target genes, we obtained fine-mapped eQTL data from the GTEx v8 release matching the human haplotype measurements (GTEx_v8_finemapping_DAPG.txt,^27^). This dataset contains 21,648,584 fine-mapped eQTLs across 49 tissues. We extracted positional information and target gene names, resulting in a subset of 21,412,255 eQTLs across tissues, comprising 2,740,212 unique eQTLs. We then overlapped the positions of the eQTLs with the linked ncRNA loci and classified a linkage as confirmed if an overlapping eQTL was predicted to regulate the same target gene as predicted for the ncRNA.

### Intersecting GWAS-SNPs with linked ncRNAs

To elucidate the functional roles of non-coding GWAS variants, we downloaded all associations v1.0 from the NHGRI-EBI GWAS Catalog (gwas_catalog_v1.0-associations_e110_r2023-10-11.tsv,^44^). This dataset comprised the SNP information for 132,201 unique variants. After the removal of SNP x SNP interactions and non-mappable variants, we maintained 119,287 SNPs. We then overlapped these SNPs with the filtered informative ncRNA loci from the GENCODE v26 annotation (*n* = 1,825), resulting in 1,059 variants. Subsequently, we intersected the data with the linked ncRNAs (*n* = 324) to map risk variants to protein-coding targets.

## Supplementary Figures

**Supplementary Figure S1.**
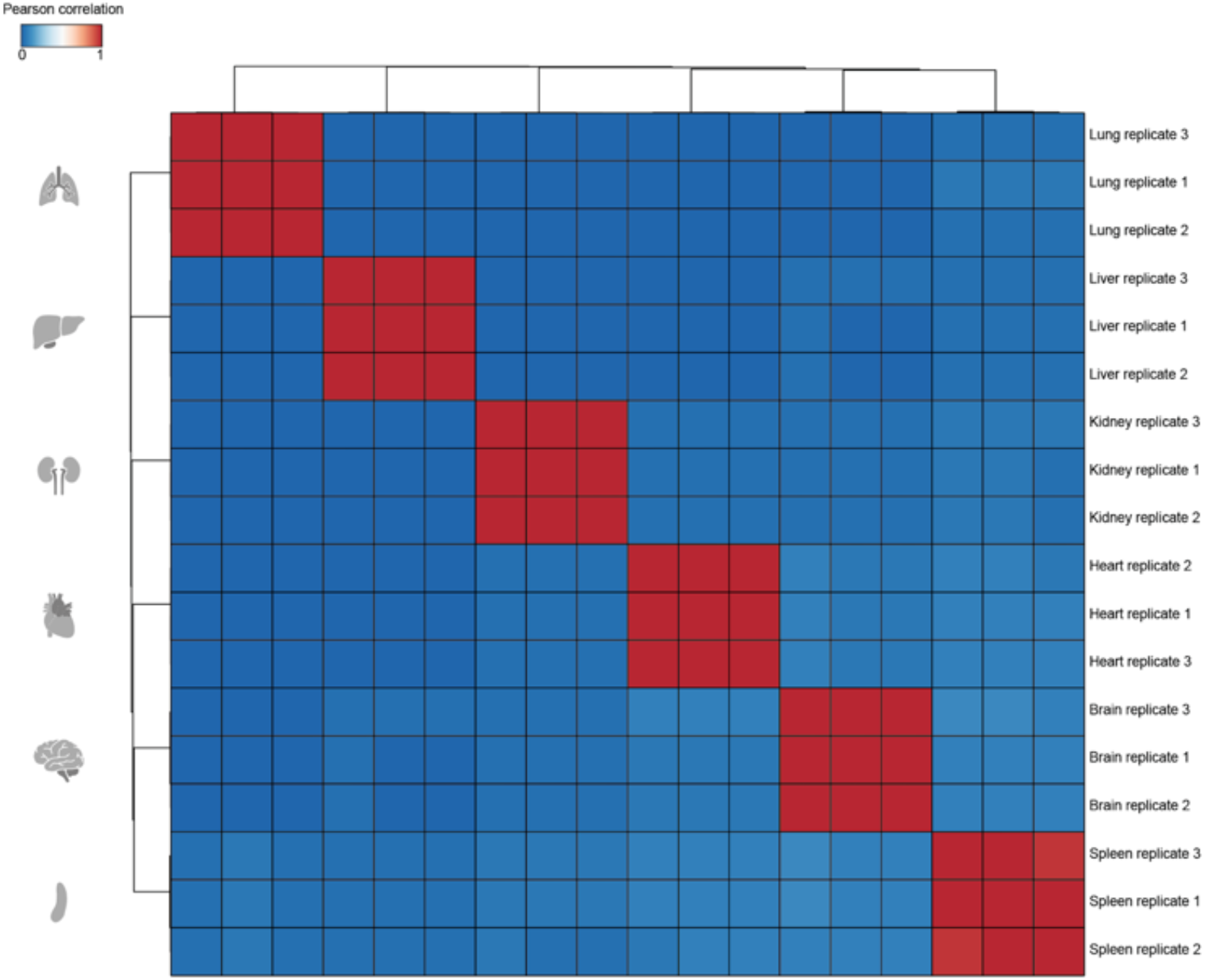
Pearson correlation matrix to confirm sample clustering. Heatmap showing unsupervised clustering of the Pearson correlation calculated on TPM values for adult tissue samples from lung, liver, kidney, heart, brain and spleen samples (*n* = 18).

**Supplementary Figure S2.**
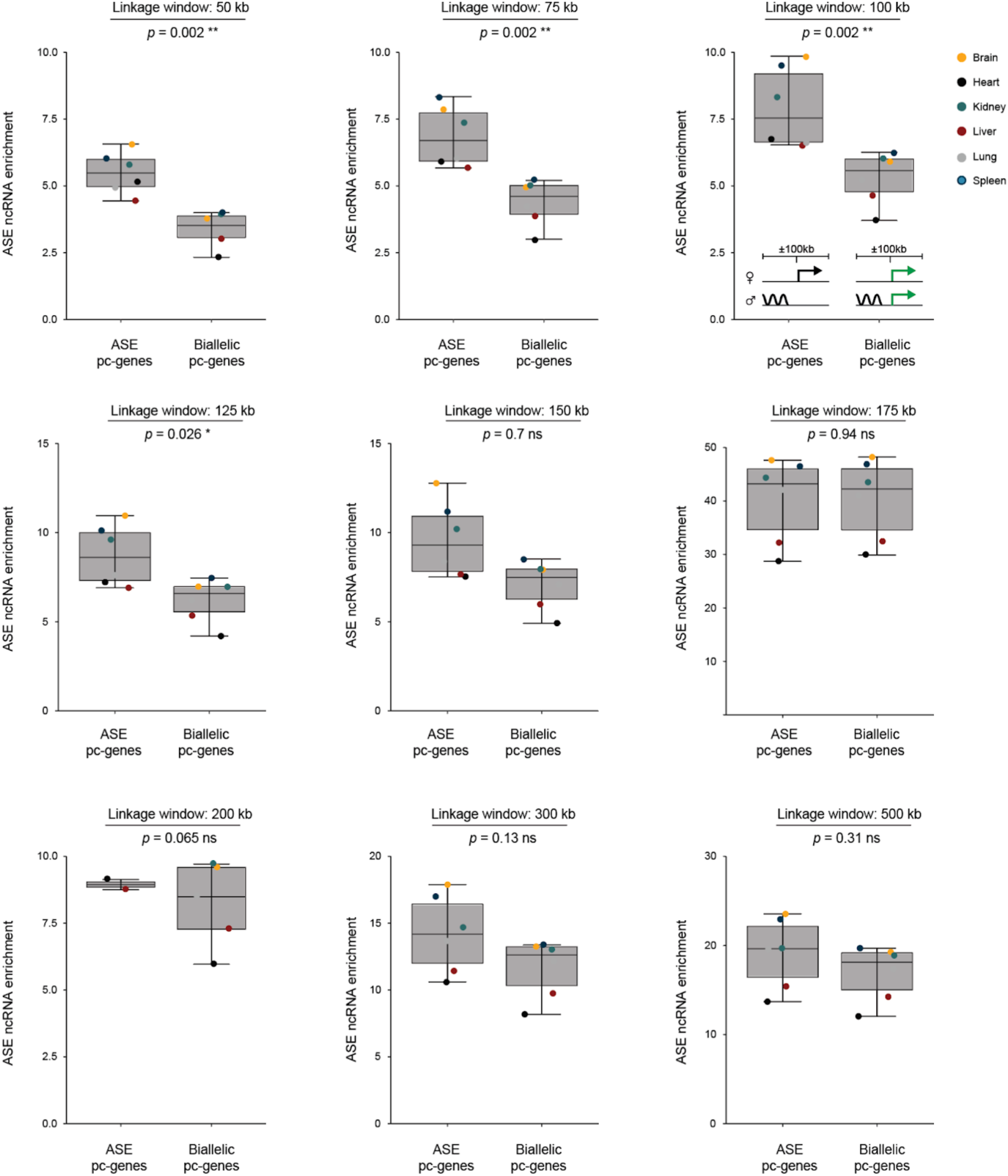
Enrichment of allele-specific ncRNAs nearby protein-coding genes across different window sizes. Box plots illustrating the percentages of allele-specific ncRNAs nearby allele-specific and biallelic protein-coding genes for various window sizes ranging from 50 - 500Kb. Boundaries of boxes indicate the interquartile range around the median. Whiskers range from maximum to minimum values. Wilcoxon tests were used to compare the allele-specific ncRNA enrichment around allele-specific and biallelic genes (*p* = 0.002). Asterisks indicates the significance level. Pc: protein-coding, ns: non-significant.

**Supplementary Figure S3.**
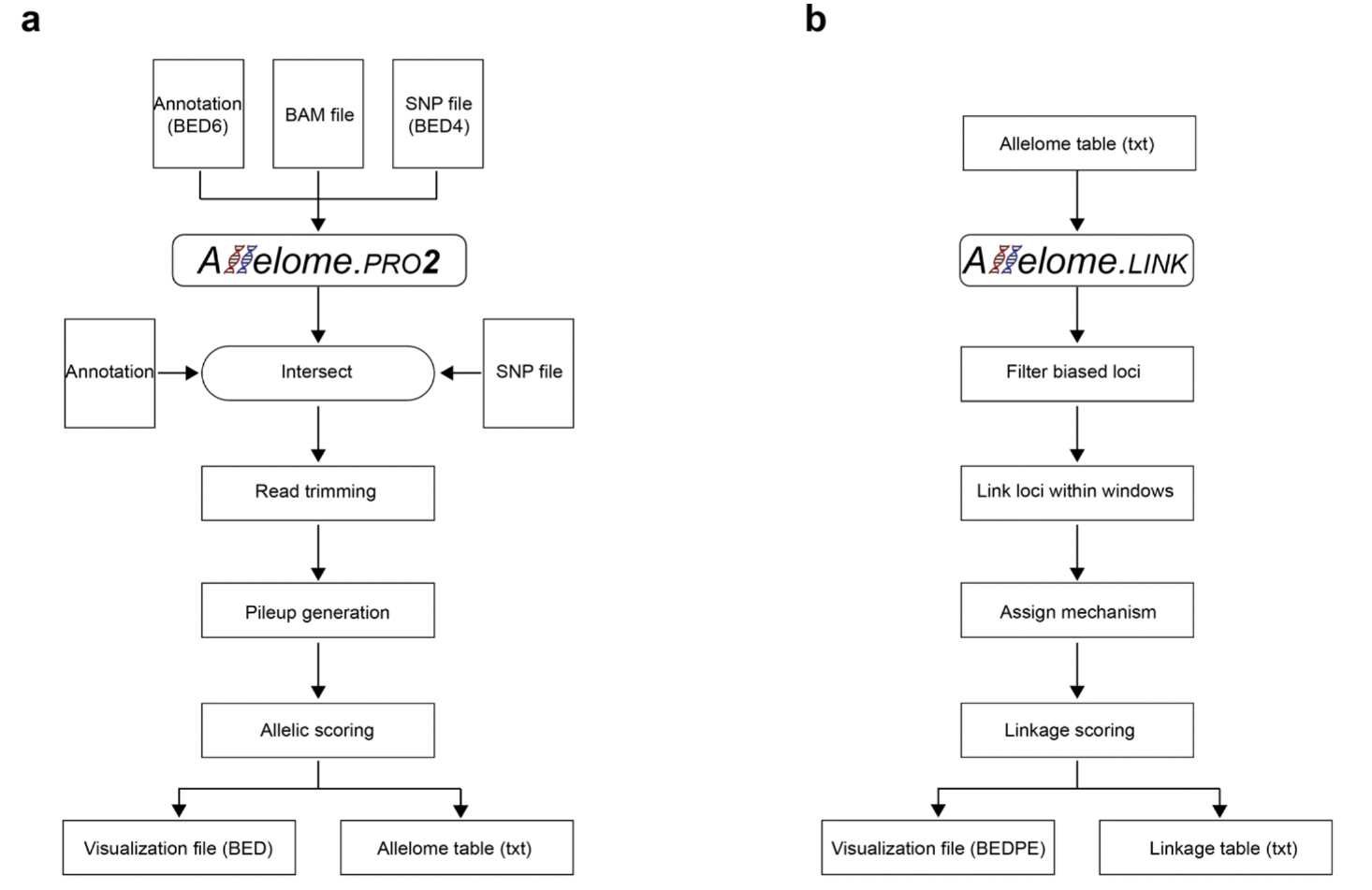
Overview of the Allelome.PRO v2.0 and Allelome.LINK pipeline. (**A**) Overview of the Allelome.PRO v2.0 pipeline. Allelome.PRO v2.0 requires three input files: an annotation file, a BAM file, and a SNP file. Allelome.PRO2, first intersects the annotation with the SNP file. Then, read trimming is performed to ensure that each read overlaps only one SNP. Sequencing reads with their corresponding SNPs are stored in a pileup file, which is used for allelic scoring. As output, Allelome.PRO2 generates a visualization file and a table containing the allelic ratio profiles of the sequencing reads. (**B**) Overview of the Allelome.LINK pipeline. Allelome.LINK uses the locus_table.txt output from Allelome.PRO2 as input. Biased loci are filtered according to a specified allelic ratio cut-off. Loci with sufficient read coverage are then linked within predefined window sizes. The mechanism for these linkages is determined based on allelic correlation or anti-correlation. A linkage score is calculated for each identified interaction. The output includes a visualization file and a linkage table containing the predicted linkages.

**Supplementary Figure S4.**
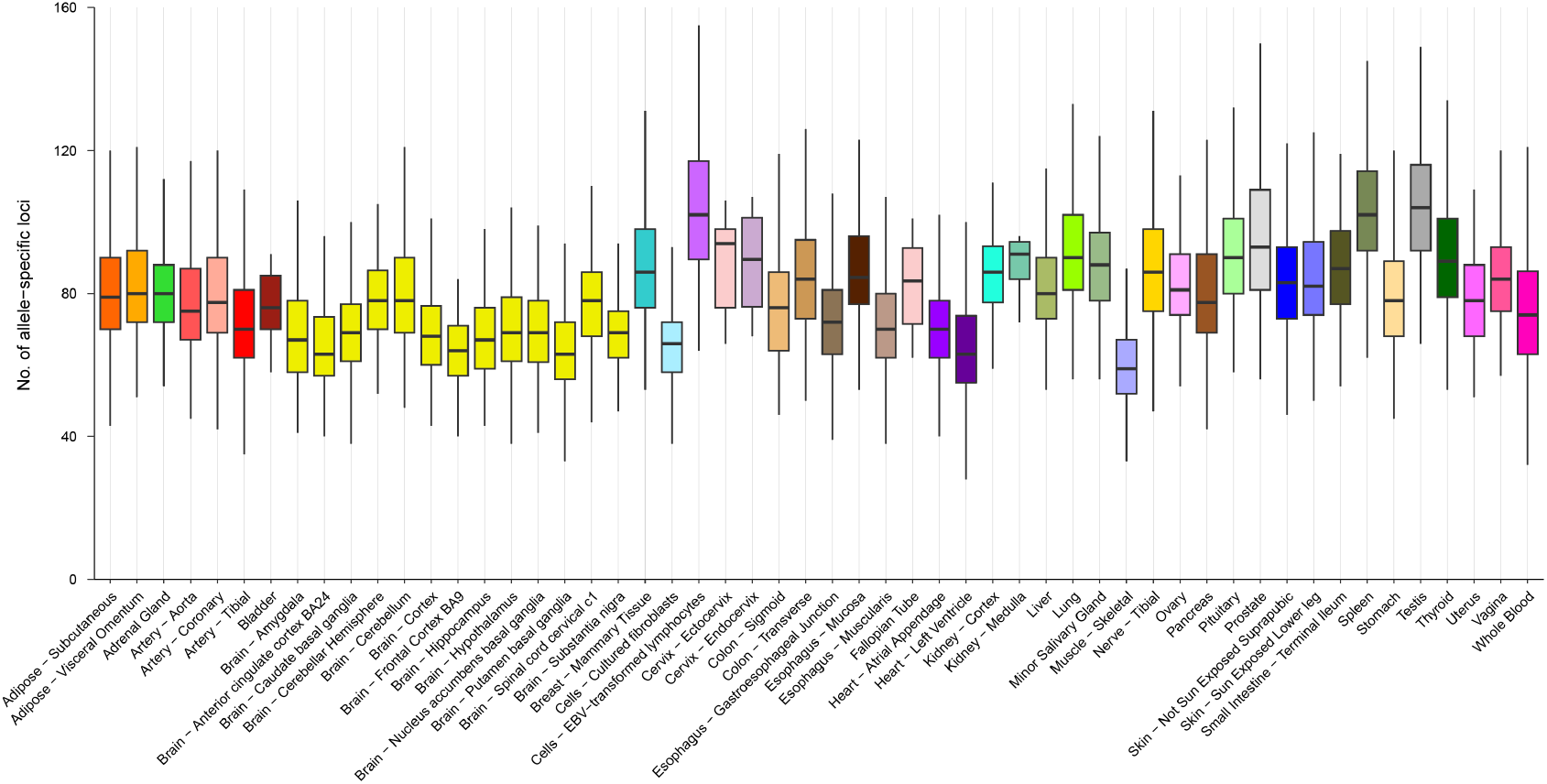
Number of allele-specific loci for individuals across 54 human tissues. Boxplots show the number of allele-specific loci for individuals from each of the 54 different human tissues obtained from the GTEx database. Boundaries of boxes indicate the interquartile range around the median. Whiskers range from maximum to minimum values. Colors represent the different tissue samples.

